# Evo 2′s Perception of Single Nucleotide Substitutions in the Genes of Two Plant Model Organisms

**DOI:** 10.64898/2026.07.01.729829

**Authors:** Otho Mantegazza, Lorenzo Bertolini, Gabriele Leoni, Moreno Colaiacovo, Mauro Petrillo, Laura Bonfini, Cristian Savini, Mario Ceresa, Xavier Zaoui

## Abstract

Although DNA Large Language Models (DNA-LLMs) offer a path to decoding genetic complexity, our ability to evaluate these models is constrained by our incomplete understanding of the very same genetic syntax and functional logic that these models are trained to learn. In this study we use single nucleotide substitutions that have or have not been observed in living organisms, to evaluate how the DNA-LLM Evo 2 interprets gene sequences from two plant model organisms, *Arabidopsis thaliana* and *Oryza sativa japonica*. Using perplexity as a measure of the model’s confidence, we observe that alleles containing simulated substitutions are perceived, on average, as less likely than those observed *in vivo*. Although the size of the effect is modest, the effect is statistically significant and robust, suggesting that Evo 2 is aligned with our current understanding of evolutionary selective constraints. This approach is designed to be model-agnostic and species-agnostic and could serve as a generic framework for evaluating the performance of DNA-LLMs.

## 1 Introduction

While the adoption of foundation large language models (LLMs) has become ubiquitous (Maslej et al., 2025), evaluating their performance across diverse professional domains is an increasingly complex undertaking. Currently, foundation LLMs are expected to match the proficiency of skilled professionals, including comprehension, manipulation and generation of natural language across diverse knowledge domains. Consequently, foundation models are evaluated using frameworks that resemble cognitive tests and reasoning-based assessments typically required for skilled professional roles. For a pragmatic overview of these evaluation practices, see Rudd et al. (2025), and Chapter 2 of Maslej et al. (2025).

DNA sequences are frequently represented as text strings because their structure and function resemble those of natural written languages (Searls, 2002). Like written languages, DNA sequences are linear, using a four nucleotides “alphabet” where the specific sequence encodes complex information. Moreover, in protein-coding regions the mapping of codons to amino acids resembles the relationship between graphemes and phonemes in linguistics (Martindale et al., 1996).

For these reasons, architectures and training procedures used in LLMs are being adapted for the challenge of interpreting text-based representations of DNA sequences. Then, models are being trained on vast amounts of genetic information, resulting in foundation DNA Large Language Models (DNA-LLMs), according to the definition for foundation models provided by Bommasani et al. (2022). These models could enable the analysis of DNA syntax and grammar, facilitating tasks such as predicting the effect of mutations, optimizing sequences across diverse species, and generating novel functional sequences for bioengineering and biomanufacturing. Examples of such models are: Nucleotide Transformer (Dalla-Torre et al., 2024), DNABERT-2 (Zhou et al., 2024), Evo 2 (Brixi et al., 2026), and, recently, the multimodal model AlphaGenome (Avsec et al., 2026). These models are already being used, for example, to engineer *ex-novo* simple viral genomes (King et al., 2025).

DNA sequences fulfill at least three intertwined roles: (i) encoding genetic information in both protein-coding and non-coding sequences, (ii) providing a structural scaffold for that information, and (iii) serving as substrate for evolutionary change. Indeed, genomes contain highly functional sequences interspersed with structural elements and with regions that appear non-functional; these parts are often so tightly integrated that they lack clear boundaries. These functional and structural regions can be identified by their evolutionary conservation (Haerty & Ponting, 2014), as exemplified by the case of long non-coding RNAs (Ponting & Haerty, 2022). Conversely, non-functional parts, which mutate faster, are likely to offer long-term evolutionary advantages by serving as a reservoir for future innovation (Van Oss & Carvunis, 2019). In plants, the most functional parts of the genome exhibit lower mutation rates due to the combined effect of evolutionary pressure and targeted DNA repair mechanisms, as reviewed in Quiroz et al. (2023).

Despite the constant and significant advances in our understanding of genomic syntax and of evolutionary dynamics, the evaluation of foundation DNA-LLMs faces a challenge that is arguably unique to the biological domain. Unlike natural languages or programming languages, DNA is a product of biological evolution rather than of intentional communication. This distinction suggests that human cognition may lack a native template for the intuitive processing of genetic sequences. While Ivanova et al. (2020) demonstrated that even human-made programming languages fail to activate the brain’s linguistic centers, considering also the findings on the cognitive constraints on natural language reported by Futrell & Hahn (2025), we expect this effect to be even more severe for DNA, which displays unique linguistic patterns and unequal entropy across regions (Akhmetov et al., 2025).

This situation creates a concurrence of two distinct interpretability challenges: researchers are tasked with evaluating a complex model against a similarly enigmatic biological code. Thus, DNA-LLM evaluation cannot easily draw from nuanced cognitively-inspired benchmarks used for other foundation LLMs. Instead, the field may rely only on frameworks grounded on our empirical knowledge of evolution, and of functional and structural genetics.

In this study, we use single nucleotide substitutions, arguably the simplest form of mutation, to study how DNA-LLMs interpret DNA sequences of protein-coding genes, by measuring which class of mutation makes the model more perplexed. Because of our limited mechanistic understanding of both the model internal representations and of the information encoded in DNA, we rely on random sampling to make minimal assumptions about the single nucleotide substitutions used in our investigation. The sampled mutations are classified according to two fundamental categories based on (i) functional annotation and on (ii) whether the mutation has been reported *in vivo* or simulated computationally. Regarding model selection, we focus on the Evo 2 model (Brixi et al., 2026), currently the open source DNA-LLM with the broadest training set and single-nucleotide resolution (Balakrishnan et al., 2025; Larey et al., 2026), which scores the highest in current benchmarking efforts (Balakrishnan et al., 2025; Larey et al., 2026), using it in its 7 billion parameters version (Evo 2 7B), optimal for genesized sequences (Brixi et al., 2026). As for the source of DNA sequences, we decided to focus on gene sequences from two plant species whose genomes have been extensively described: *Arabidopsis thaliana*, a fundamental model organism for plants, and *Oryza sativa japonica*, model organism for cereal crops.

## 2 Material and Methods

### 2.1 Technology stack

This study is written reproducibly in literate programming style; source code and DNA sequences are available at https://data.europa.eu/doi/10.2905/JRC.JH9JHTG. All analytic steps besides Evo 2 inference, are streamlined and can be run on standard consumer-grade hardware.

Evo 2 7B inference was run on NVIDIA H100 80GB GPUs with a workflow documented in Section 2.4.

Any other analyses were run with R version 4.6.0 (R Core Team, 2025). The full stack of R packages used is recorded with RENV version 1.0.11 (Ushey & Wickham, 2026), bioinformatics operations were run with Biocon-ductor version 3.23 (Huber et al., 2015), data operations were run with Tidyverse version 2.0.0 (Wickham et al., 2019). The whole manuscript can be regenerated with Quarto, version 1.9.16 (Allaire et al., 2025).

### 2.2 Data source

All genomic data, including genomic sequences, functional annotations and natural variations were sourced from Ensembl Plants version 60, which is part of Ensembl Genomes (Yates et al., 2021).

Ensembl Plants fulfilled our requirements for a source of genomic information that (i) spans multiple species through a unified interface, (ii) is versioned and (iii) is openly accessible. These aspects are crucial for evaluating DNA-LLM’s comprehension of DNA sequences. We especially value these properties of Ensembl:

- Unified access to coordinated genomic data, including reference genomic sequences, structural and functional annotations and known natural variation.
- Ease of programmatic access through FTP, HTTP and REST API.
- Explicit versioning of both the tool and the data, while keeping previous versions accessible (we used version 60).
- Long lasting commitment to reliable public infrastructure: the Ensembl project was launched in 1999 (Hubbard et al., 2002) focusing on the human genome; then including animal model organisms. The project was further expanded in 2009 by launching Ensembl Genomes (Yates et al., 2021), which extends this infrastructure to non-vertebrates. Today it covers bacteria, protists, fungi, metazoa and plants.
- Commitment to free circulation of information: all data are licensed openly under no restriction, software is under the open source license Apache 2.0.

### 2.3 Sequence acquisition, selection and substitution

The preprocessing workflow, encompassing (i) sequence and annotation acquisition from Ensembl Plants, (ii) gene selection, and (iii) nucleotide substitution, was implemented via a suite of R scripts (see Section 5) and executed through the `R/preprocess.R` wrapper script.

To ensure high-confidence genomic annotations across species, we initially selected genes from the Ensembl Plants database containing valid `gene_name` entries. We extracted a random sample of 500 genes for both *Arabidopsis thaliana* and *Oryza sativa japonica*, however 68 were subsequently excluded from the *Arabidopsis thaliana* gene pool, as they were identified as non-coding RNAs (ncRNAs), which fall outside the scope of this study.

After extracting the genomic sequence for each gene —defined as the region from the earliest transcription start site (TSS) to the latest transcription termination site (TTS)—two classes of single nucleotide substitutions were introduced to the filtered sequence pool: (i) all known variants reported in the Ensembl Plants variation database; and (ii) synthetic mutations introduced at a random 5% of nucleotide positions. For the latter, each reference nucleotide was substituted by all three possible alternative alleles, generating three distinct mutant sequences per targeted site.

### 2.4 Evo 2 7B inference

To run Evo 2 7B inference, we used the official `Evo2` Python library provided by the authors, with default configuration and hyperparameters (Brixi et al., 2026). The raw output logits were transformed into the model-assigned next-nucleotide probabilities using the softmax function, as implemented in the `Pytorch` framework. To optimize storage and downstream computational efficiency, the probability estimates were cast to 16-bit-floating point precision (`float16`) and exported as compressed numpy arrays.

The probability estimates were imported into R using the `reticulate` package, version 1.46.0 (Ushey et al., 2025). We developed custom scripts to extract next-nucleotide predictions, which were executed through the `R/preprocess.R` wrapper.

To assess the predictive proficiency of the model over a whole sequence, we opted to use perplexity (PPL) as intrinsic evaluation metric (Miaschi et al., 2021).

For a given sequence of length *n* nucleotides, perplexity (*P P L*) is the exponentiated average negative log-likelihood of a sequence, calculated as:

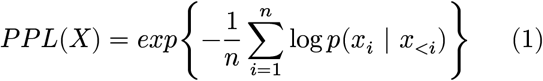

where log *p*(*x*_*i*_ | *x*_<*i*_) is the model-assigned probability of the observed nucleotide *x* given the preceding sequence *x*_<*i*_.

Following the approach of scoring both forward and reverse-complement strands for a given nucleotide sequence, described by Brixi et al. (2026), we calculated perplexity independently for both the forward and reverse-complement strands. The final reported values represent the arithmetic mean of these two estimates.

Lower perplexity values indicate a more “certain” model that is less surprised by the sequence under evaluation.

### 2.5 Data Analysis

To describe and analyze the next nucleotide predictions and perplexities obtained from Evo 2 7B inference, we favored robust statistics, nonparametric tests and extensive exploration of results via data visualization and tables. All these steps were implemented in R 4.6.0 (R Core Team, 2025).

To measure association we calculated Spearman’s rank correlation coefficient (ρ). To compare overall distribution shapes between groups, we employed the two-sample Kolmogorov-Smirnov asymptotic test. Differences in central tendencies between two groups were evaluated using the Wilcoxon Rank-Sum test, for which, beyond the p-value we reported the probability of superiority (Fiel Peres, 2026; Vargha et al., 2000) as an effect size. This effect size metric corresponds to the area under the receiver operating characteristic curve (AUC). The *AUC* was estimated from the Mann–Whitney *U* statistic as:

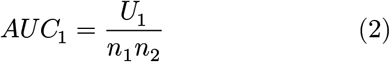

where *n*_1_ and *n*_2_ represent the sample sizes of the respective groups. All tests were conducted as implemented in the R `stats` package, which is natively part of R.

To estimate the proportion of perplexity variance that can be attributed to differences between genes, we calculated the intraclass correlation coefficient (Nakagawa et al., 2017). A linear mixed effect model was fitted with `perplexity` as outcome and `gene ID` as random intercepts, using the `lme4` package version 2.0-1 (Bates et al., 2015). The intraclass correlation coefficient was then extracted from the mixed effect model using the `performance` R package, version 0.16.0 (Lüdecke et al., 2021).

Bootstrapped confidence intervals were estimated with 1000 replicates using the `rsample` package, version 1.3.2.

All visualizations were produced with `ggplot2` version 4.0.3 (Wickham, 2016) and all tables were produced with `gt` version 1.3.0 (Iannone et al., 2026).

## 3 Results and Discussion

### 3.1 Selection of the Experimental Setup for Single-Nucleotide Substitution

In order to standardize our analysis across genomic loci, we decided to focus on protein-coding genes. We ran inference with Evo 2 7B on gene sequences extracted as they are from the reference genome, and on the same sequences modified by changing exclusively one single nucleotide. To extract gene sequences, we relied on the functional annotations of the reference genomes of *Arabidopsis thaliana*, version TAIR10 and *Oryza sativa japonica*, version IRGSP-1.0, as reported in Plant Ensembl 60, spanning from the 5′ UTR of the earliest-starting isoform to the 3′ UTR of the latest-ending isoform.

Analyzing the entire potential set of observed and simulated single-nucleotide substitutions across the full set of genes of *Arabidopsis thaliana* and *Oryza sativa japonica*, would be computationally prohibitive. To reduce computation time, we sampled pseudorandomly 500 genes per species which, after retaining protein-coding genes, resulted in 432 genes out of 27628 for *Arabidopsis thaliana* and 499 out of 35804 for *Oryza sativa japonica*.

Within these genes, single nucleotide substitutions were applied to a 5% pseudorandom sample of all possible positions. Additionally, we incorporated all single nucleotide variations observed *in vivo* and recorded in the Ensemble Plants variation database, yielding a total of 351894 substituted sequences for *Arabidopsis thaliana* and 468984 for *Oryza sativa japonica*. These substituted sequences, together with their respective unmodified sequences, were analyzed with Evo 2.

### 3.2 Description of the substituted sequences

Mutations were classified into four functional biological categories based on gene annotations. Those located in non-coding regions (UTRs and introns) were labeled as (i) non-coding, while those within protein-coding regions, were categorized according to their effect on the codon as: (ii) silent, (iii) missense, or (iv) nonsense. These four functional classes are combined with one variable that indicates whether the mutation is recorded in Ensembl’s variant database. Accordingly, mutations are classified as either (i) Observed, if recorded, or (ii) Simulated, if not. The frequencies with which these combined categories appear in the dataset developed for this study are reported in Table 1. Since simulated mutations are introduced randomly at a uniform rate, the ratio of observed mutations over the total mutation pool (observed and simulated) serves as a proxy for the frequency at which mutations are observed and recorded across a given genomic section.

**Table 1:** Number of single nucleotide mutations category. Observed mutations are sourced from Ensembl 60 VCF files, while simulated mutations are introduced randomly at a uniform rate. **Ratio Observed [%]**, the ratio of observed mutations over the total mutations (observed + simulated) serves as a proxy for recording frequency in the Ensembl database. The ratio varies consistently across both species: silent mutations show the highest relative rate, followed by non-coding, missense, and nonsense mutations. All mutation types were observed at a higher ratio in *Arabidopsis thaliana* than in *Oryza sativa japonica*.

| Mutation Type | Observed [n] | Simulated [n] | Ratio Observed [%] |
| --- | --- | --- | --- |
| *Arabidopsis thaliana* |  |  |  |
| Mutation in non-coding region | 42 364 | 146 240 | 22.46% |
| Silent mutation | 12 795 | 32 171 | 28.45% |
| Missense mutation | 18 963 | 92 779 | 16.97% |
| Nonsense mutation | 472 | 6 110 | 7.17% |
| *Oryza sativa japonica* |  |  |  |
| Mutation in non-coding region | 52 190 | 269 005 | 16.25% |
| Silent mutation | 7 780 | 33 269 | 18.95% |
| Missense mutation | 10 427 | 91 365 | 10.24% |
| Nonsense mutation | 262 | 4 686 | 5.30% |

Across both species, silent mutations are observed and recorded in public databases at the highest rate, followed by mutations in non-coding regions, missense and, last, nonsense mutations (Table 1). These varying rates among the four functional classes reflect the natural biases of the process of mutation, repair and selection, reviewed in Quiroz et al. (2023).

The higher average rate with which mutations are observed and recorded in *Arabidopsis thaliana* compared to *Oryza sativa japonica* could, instead, be ascribed to a more extensive characterization of the former’s genome. Indeed, the category of observed mutations represents only a sample of those likely to appear in nature, which might randomly fall in the simulated category.

### 3.3 Evo 2 accurately predicts the next nucleotide for protein-coding sequences

We initially described Evo 2 7B next-token (i.e. next-nucleotide) prediction performance on unmodified gene sequences from the reference genomes. These results are shown as the model’s assigned probability for the correct next nucleotide, estimated using the preceding sequence as context, starting from the gene’s earliest transcription start site to the nucleotide immediately prior to the target.

As shown in Figure 1, Evo 2 7B predicts protein-coding sequences with a probability approaching 1. In contrast, predictions for non-coding sequences center near 0.25—a value equivalent to random selection among four nucleotides (A, T, C and G). This pattern is consistently observed across representative examples: gene AT1G03190 from *Arabidopsis thaliana* and gene Os09g0567366 from *Oryza sativa japonica*.

**Figure 1:**
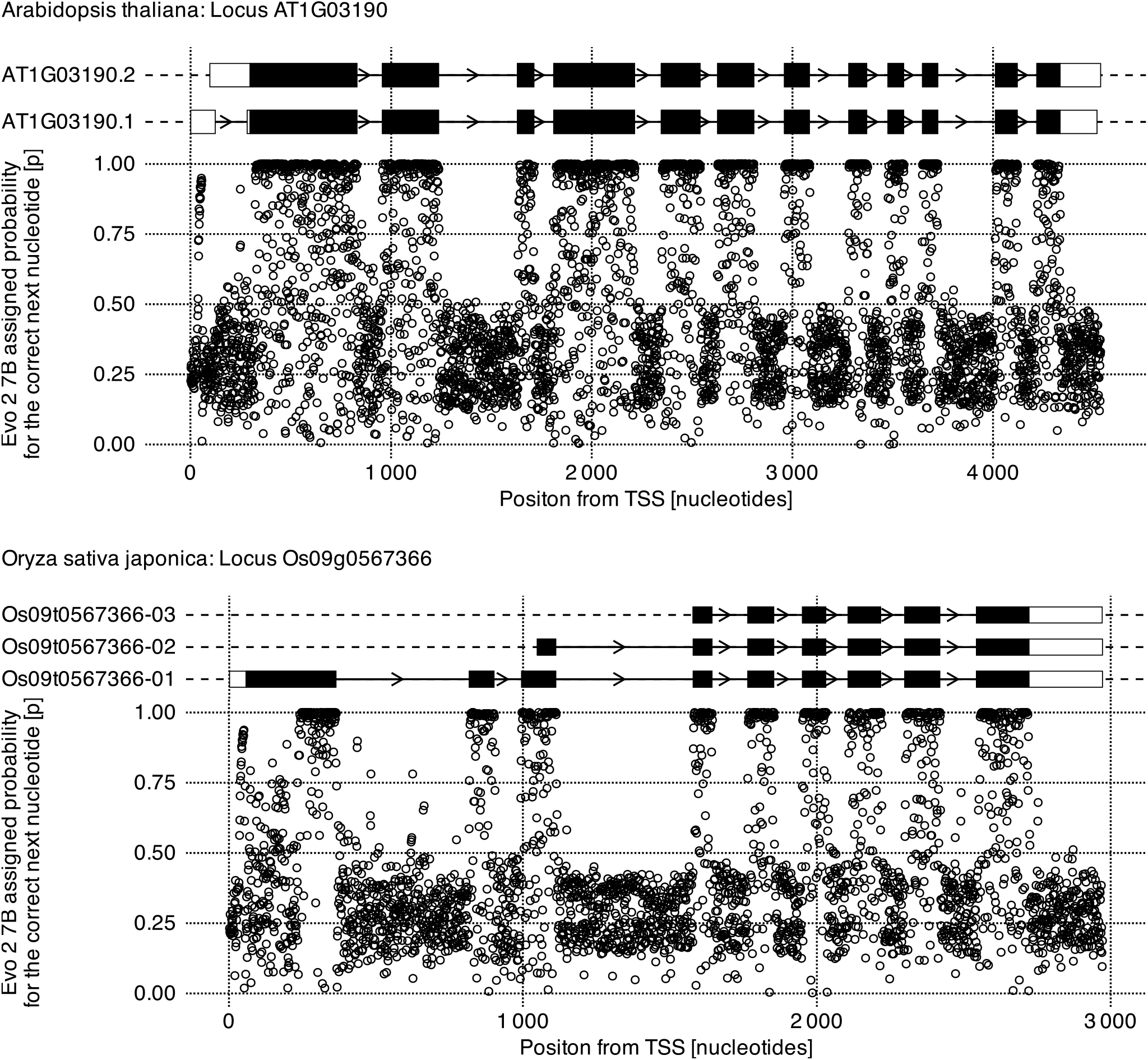
Next nucleotide predictions along gene sequences. Evo 2 7B assigned probability for the correct next nucleotide along two representative genes: *Arabidopsis thaliana* gene AT1G03190 and *Oryza sativa japonica* gene Os09g0567366. These examples illustrate the model’s predictive performance on unmodified genomic sequences. In each panel, dot plots at the bottom represent Evo 2′s probabilities, while genomic features are represented at the top as follows: black boxes for protein-coding sequences, white boxes for exonic non-coding sequences (UTRs) and black arrowed lines for introns; distinct lines represent known splicing isoforms. Evo 2 predicts protein-coding sequences with high confidence.

Figure 2 aggregates Evo 2 7B predictions for all unmodified genes in this study, confirming the pattern described above, while highlighting notable deviations. While the primary peak approaches 1 for protein-coding sequences and 0.25 for non-coding sequences, minor reciprocal peaks are also evident; thus, a minor peak centered close to 1 is clearly visible in non-coding sequences, and, likewise, a minor peak centered at 0.25 is visible in coding sequences.

**Figure 2:**
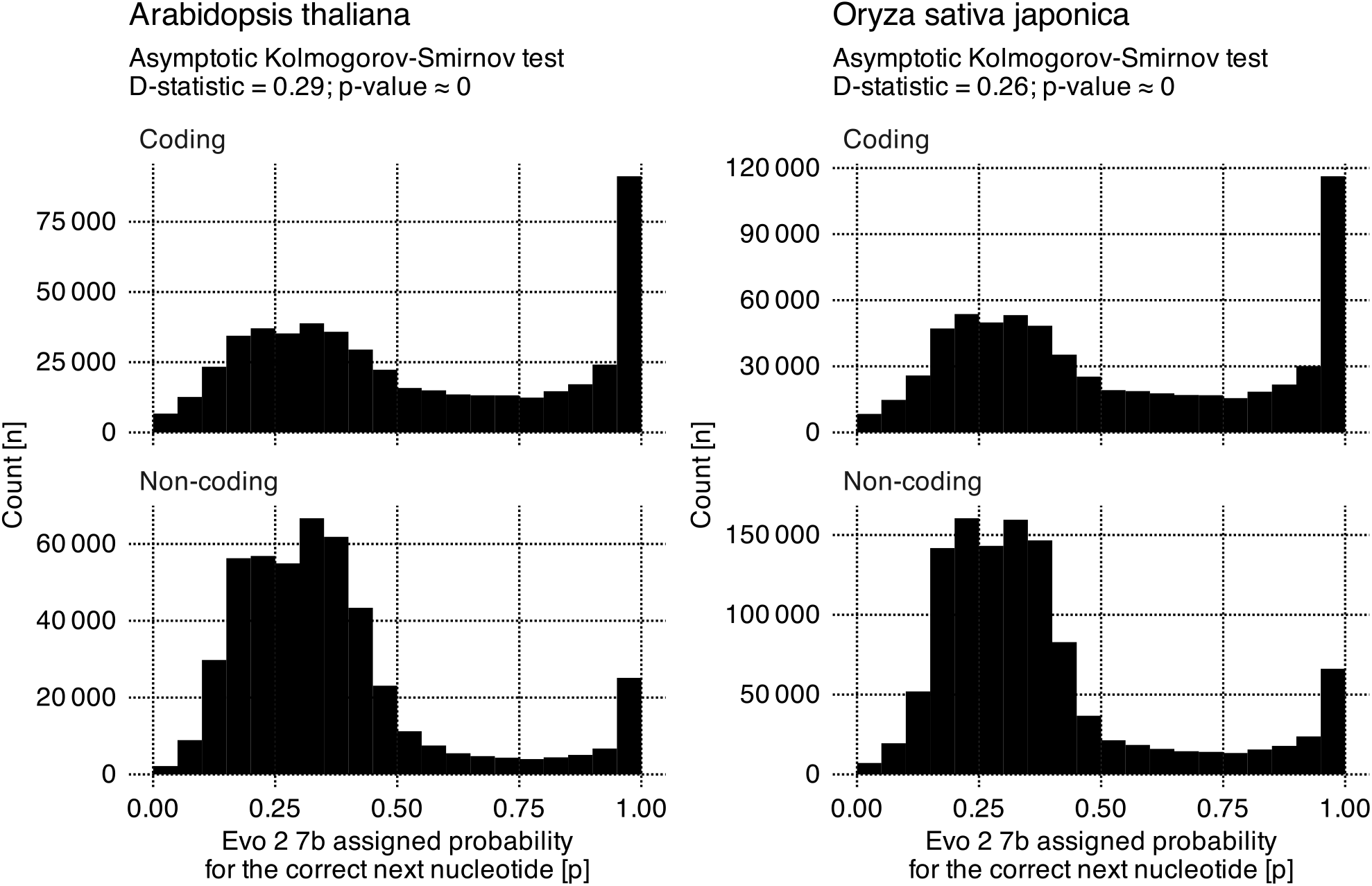
Next nucleotide predictions for protein-coding and non-coding sequences. Histograms display Evo 2 7B assigned probability for the correct next nucleotide, across all unmodified gene sequences considered in this study. In both *Arabidopsis thaliana* and *Oryza sativa japonica*, probabilities for protein-coding sequences peak near 1. In contrast, non-coding sequences (UTRs and introns) show a prominent peak centered just above 0.25, which appears to be bimodal, and is characterized by two closely spaced local maxima. While a Kolmogorov-Smirnov test indicates that coding and non-coding predictions are drawn from distinct distributions, the presence of reciprocal minor peaks might indicate either sequence misannotation or model inaccuracy.

These reciprocal peaks might arise from several phenomena. While predictions might be initially hampered at the gene start—likely due to reduced sequence context —deviations also occur further downstream. In these instances, protein-coding sequences are predicted with the low probability typical of non-coding regions, and vice-versa. These discrepancies may stem from either yet-to-be described internal model mechanisms, non-coding conserved DNA sequences, such as regulatory motifs, or from misannotated genomic features, which are subjected to frequent refinement, given the inherent complexity and versatility of genomes (a process reviewed by Pilalis et al. (2025)).

Furthermore, in Figure 2, the 0.25 peaks appear bimodal, characterized by two distinct local maxima; a nuance that remains visible upon closer inspection of also the individual examples in Figure 1. The origin of the bimodal peaks in non-coding sequences is, to our knowledge, unexplained.

### 3.4 Evo 2′s perplexity for gene sequences is uncorrelated to gene length but partially correlated to gene CDS ratio

To compare the effectiveness of Evo 2 7B’s next nucleotide predictions across diverse genes, we used perplexity as the primary metric. Perplexity serves as a summary indicator of the average model uncertainty over a nucleotide sequence, consequently lower values represent a superior predictive performance over a given sequence.

We first used perplexity to investigate how the model interprets the 432 unmodified gene sequences from *Arabidopsis thaliana*, TAIR10 genome, and the 499 from *Oryza sativa japonica*, IRGSP-1.0 genome. The relationship between perplexity and fundamental genomic features: (i) gene length, and (ii) protein-coding sequence ratio (CDS ratio, calculated as the protein-coding sequence length divided by gene length) is depicted in Figure 3.

**Figure 3:**
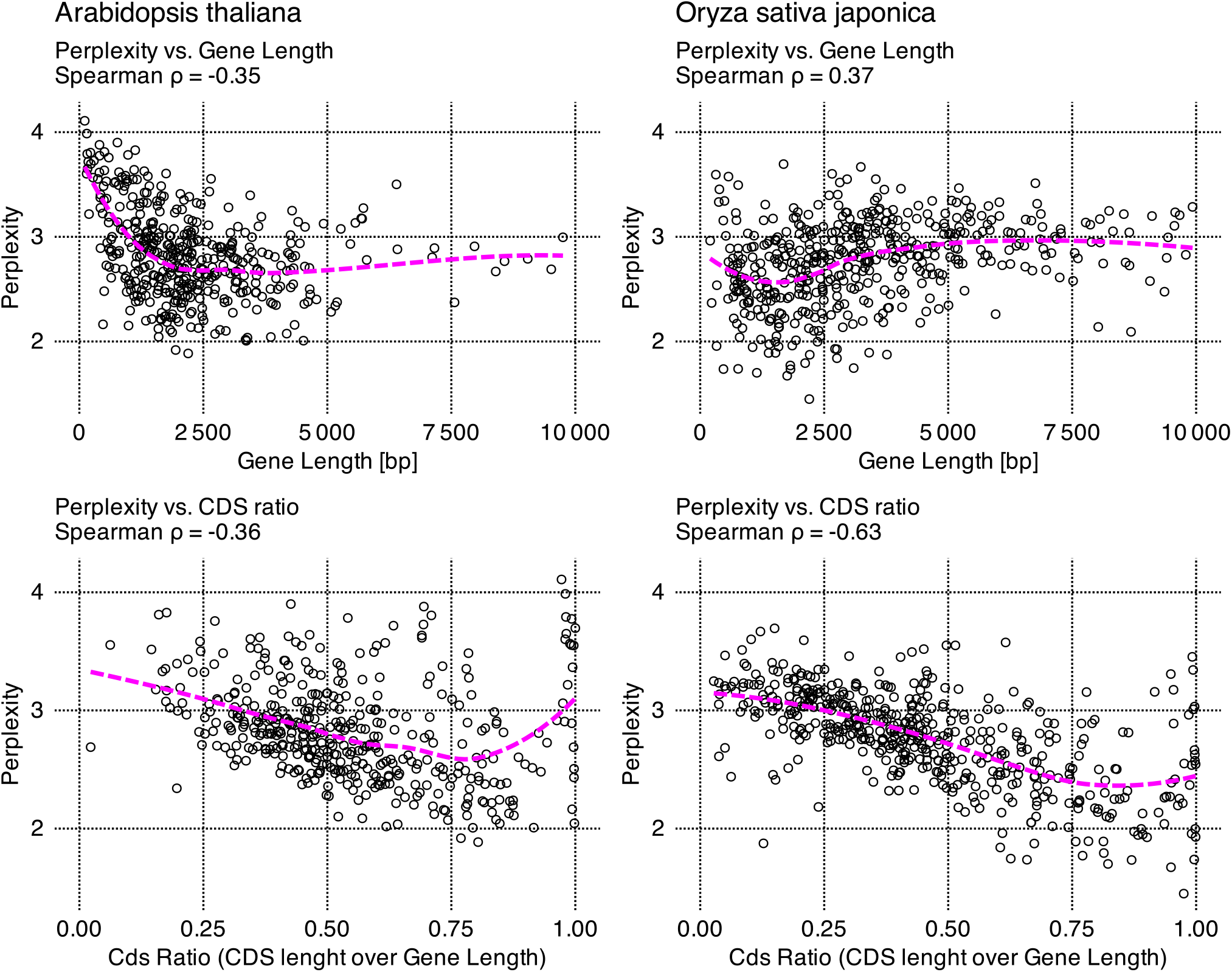
Relationship between Evo 2′s perplexities for unmodified gene sequences from reference genomes and factors such as gene length and the proportion of coding sequence (CDS ratio). In each scatterplot, individual points represent one single gene from the reference genome of *Arabidopsis thaliana*, (left column), or *Oryza sativa japonica* (right column). Across both species, the relationship between perplexity and these fundamental biological features is complex but similar. A partial correlation is evident only in the lower panels comparing perplexity and CDS ratio. To enhance readability, the x axis representing gene length is truncated at 10,000 bp, excluding 17 outlier rice genes, with exceptionally long sequences. This exclusion does not affect the findings.

As shown in Figure 3, perplexity is mostly uncorrelated with gene length (*Arabidopsis thaliana*: Spearman ρ=−0.35; *Oryza sativa japonica* ρ=0.37). However, a localized increase is observed at shorter sequences, which are more prevalent in Arabidopsis, while in rice a slightly higher average perplexity is observed for longer sequences.

In contrast, perplexity exhibits a mild negative correlation to CDS ratio in rice (ρ=−0.63). While, in Arabidopsis, the same correlation is less statistically evident (ρ=−0.36), studying the exploratory scatterplots in Figure 3 reveals a negative trend similar to the one visible in rice, interrupted only by a subgroup of genes with a localized sharp increase in perplexity at very high CDS ratios.

The independence of perplexity from gene length and its mild correlation to CDS ratio aligns with the model’s next-nucleotide prediction performance on protein-coding sequences illustrated in Figure 1 and Figure 2. Other factors influencing perplexity variations within reference genomes could be attributed to different genomic characteristics between Arabidopsis and rice, with the former being more compact, featuring fewer and shorter introns and a lower density of transposable elements (Arabidopsis Genome Initiative, 2000; International Rice Genome Sequencing Project, 2005).

Perplexity is a standard metric for assessing the capabilities of autoregressive language models (Miaschi et al., 2021) and we employ it in the following sections to investigate the effects of single-nucleotide substitutions.

### 3.5 Perplexity varies more among different genes than within single nucleotide substitutions of the same gene

After characterizing the distribution of Evo 2 perplexities for the reference alleles, we examined how single nucleotide substitutions alter those measurements. Figure 4 shows the distribution of perplexities for the substituted alleles (pooling observed and simulated variants) relative to the reference alleles.

**Figure 4:**
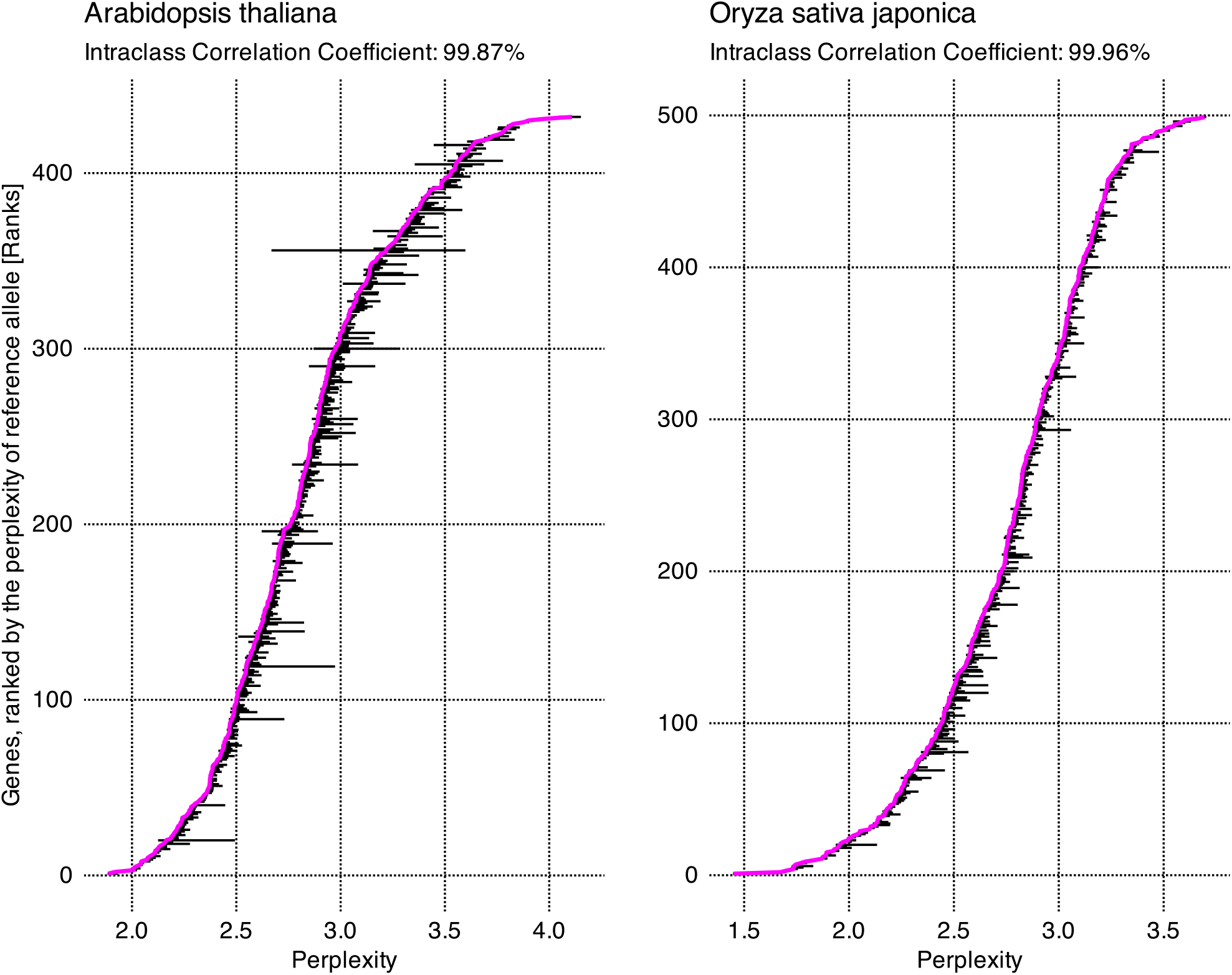
Comparative distribution of perplexity across reference and substituted alleles. Genes are ranked on the vertical axis by the perplexity of their reference allele, indicated in the magenta line. For each gene, horizontal black bars represent the central 99^th^ percentile of perplexity scores of single-nucleotide substituted alleles. In addition to the visual representation, on the top of the panels we report the intraclass correlation coefficient, indicating the proportion of variance that can be explained by the grouping structure (genes, in this case). The high intraclass correlation coefficients, together with visual evidence, indicate that perplexity variance within a single gene, is significantly lower than the overall variance across different genes. Moreover, the within-gene variance is higher in *Arabidopsis thaliana* compared to *Oryza sativa japonica*, with median IQR of 0.0045 and 0.0032 respectively.

In both species, the variance in perplexity among the substituted alleles originating from the same gene is smaller than the overall variance observed across different genes. This finding is supported by (i) visual inspection of the 99th-percentile range of perplexities for the single-nucleotide substituted alleles and (ii) the intraclass correlation coefficient, comparing within-gene variance with overall variance, that yields scores of approximately 1.0 for both species (Figure 4). Given that perplexity varies more significantly between different genes than among alleles of the same gene, all subsequent analyses were restricted to within-gene comparisons. Consequently, we restricted direct comparisons to alleles within the same gene, avoiding direct comparisons between alleles of different genes. Instead, cross-gene data are presented only as distributions of statistics derived from individual genes.

Furthermore, we sought to identify the determinants of inter-gene variation in perplexity dispersion, using the interquartile range (IQR) as our metric for within-gene spread. As illustrated in Figure 5, the magnitude of within-gene perplexity IQR correlates strongly with the gene’s overall length. (Spearman ρ of −0.90 for *Arabidopsis thaliana* and of −0.94 for *Oryza sativa japonica*).

**Figure 5:**
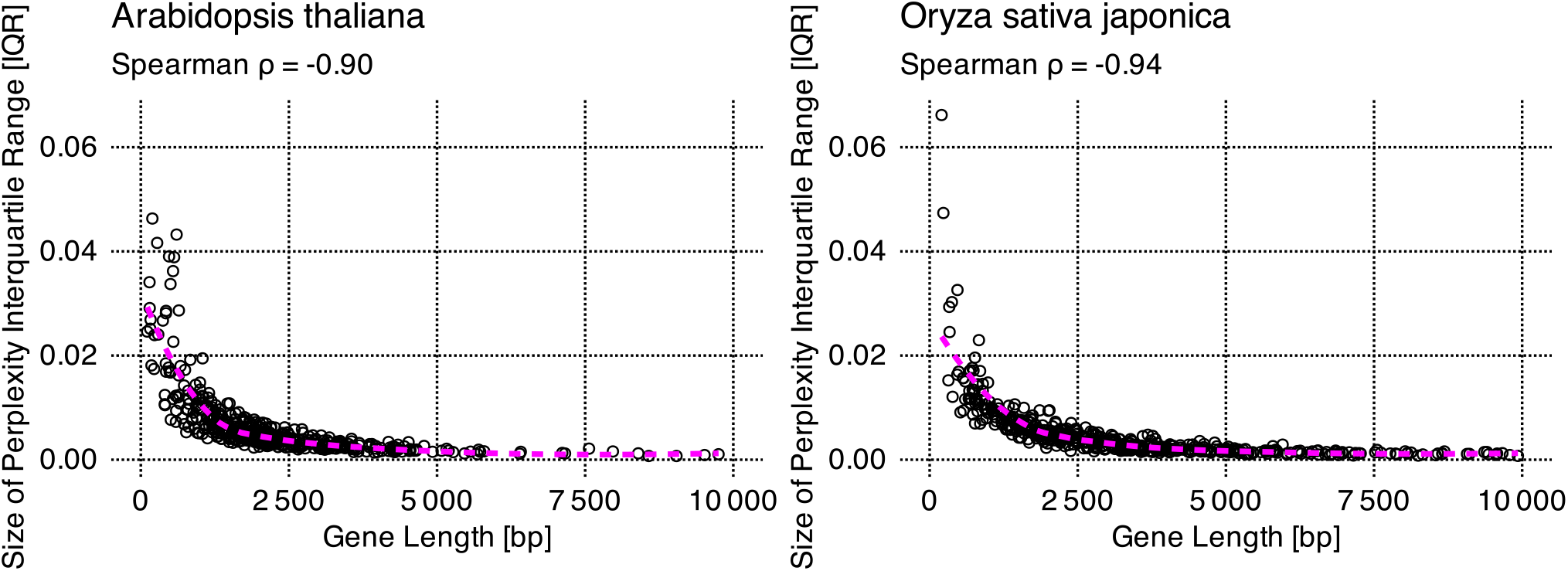
Relationship between gene length and the magnitude of the interquartile range of Evo 2′s perplexities for alleles of the same gene. In each scatterplot, a point corresponds to a single gene. The y-axis shows the spread of perplexities for that gene’s single-nucleotide-substitution alleles, quantified as the interquartile range (IQR). The IQR is strongly negatively correlated with gene length.

The attenuated impact of single-nucleotide mutations on perplexity in longer genes stems from the localized nature of Evo 2′s response. As illustrated later in this study, in Figure 9 and Table 2, the perturbation induced by single-nucleotide substitution induce on the model’s probabilities to predict the next nucleotide correctly, decays as the distance from the mutation site increases.

**Table 2:** Decay of the effect of single-nucleotide substitutions on the likelihood of downstream nucleotides. Each column reports the percentage of nucleotides downstream of the mutation whose lower-quartile likelihood difference (between substituted and reference allele) exceeds what would be expected from random noise. For clarity, distances are grouped into four orders of magnitude from the mutation (≈ 1, 10, 100, 1,000 nucleotides). In most cases the effect tends to decay toward the undetectable range within hundreds of nucleotides. The decay is slightly faster in *Oryza sativa japonica* than in *Arabidopsis thaliana*.

| Mutation Type | Strand | Distance from Mutation (Nucleotides) |  |  |  |
| --- | --- | --- | --- | --- | --- |
|  |  | [1, 10) | [10, 100) | [100, 1 000) | [1 000, 10 000) |
| Arabidopsis thaliana |  |  |  |  |  |
| Simulated | Forward | 100.0% | 100.0% | 46.6% | 0.0% |
| Simulated | Reverse-complement | 100.0% | 100.0% | 54.1% | 1.6% |
| Observed | Forward | 100.0% | 100.0% | 50.8% | 0.0% |
| Observed | Reverse-complement | 100.0% | 100.0% | 55.6% | 13.3% |
| Oryza sativa japonica |  |  |  |  |  |
| Simulated | Forward | 100.0% | 100.0% | 27.6% | 2.9% |
| Simulated | Reverse-complement | 100.0% | 100.0% | 29.2% | 4.8% |
| Observed | Forward | 100.0% | 100.0% | 28.0% | 10.0% |
| Observed | Reverse-complement | 100.0% | 100.0% | 25.1% | 7.8% |

### 3.6 Reference alleles have lower perplexity than their single-nucleotide substitution alleles

We then investigated if the perplexity of reference alleles is statistically distinguishable from that of their counterparts carrying single-nucleotide substitutions.

For each gene, we calculated the percentage of single-nucleotide substitution alleles with perplexity exceeding the one of their unmodified counterpart (shown in Figure 6). In *Arabidopsis thaliana* the median percentage of alleles with higher perplexity is 62.7% (CI with bootstrapped 2.5th–97.5th percentiles: 61.2%-64.4%) for observed mutations and 74.2% (CI: 72.6%-75.4%) for simulated mutations. In *Oryza sativa japonica* these values are 61.7% (CI: 60.0%-62.9%) and 76.8% (CI: 75.4%-78.1%) respectively. If single nucleotide substitutions had a random effect on Evo 2′s perplexity, the expected center of these distributions would be at 50%. Contrary to this expectation, we observe a distribution significantly shifted toward higher percentages across both species (*Arabidopsis thaliana* and *Oryza sativa japonica*) and both mutation classes (observed and simulated).

**Figure 6:**
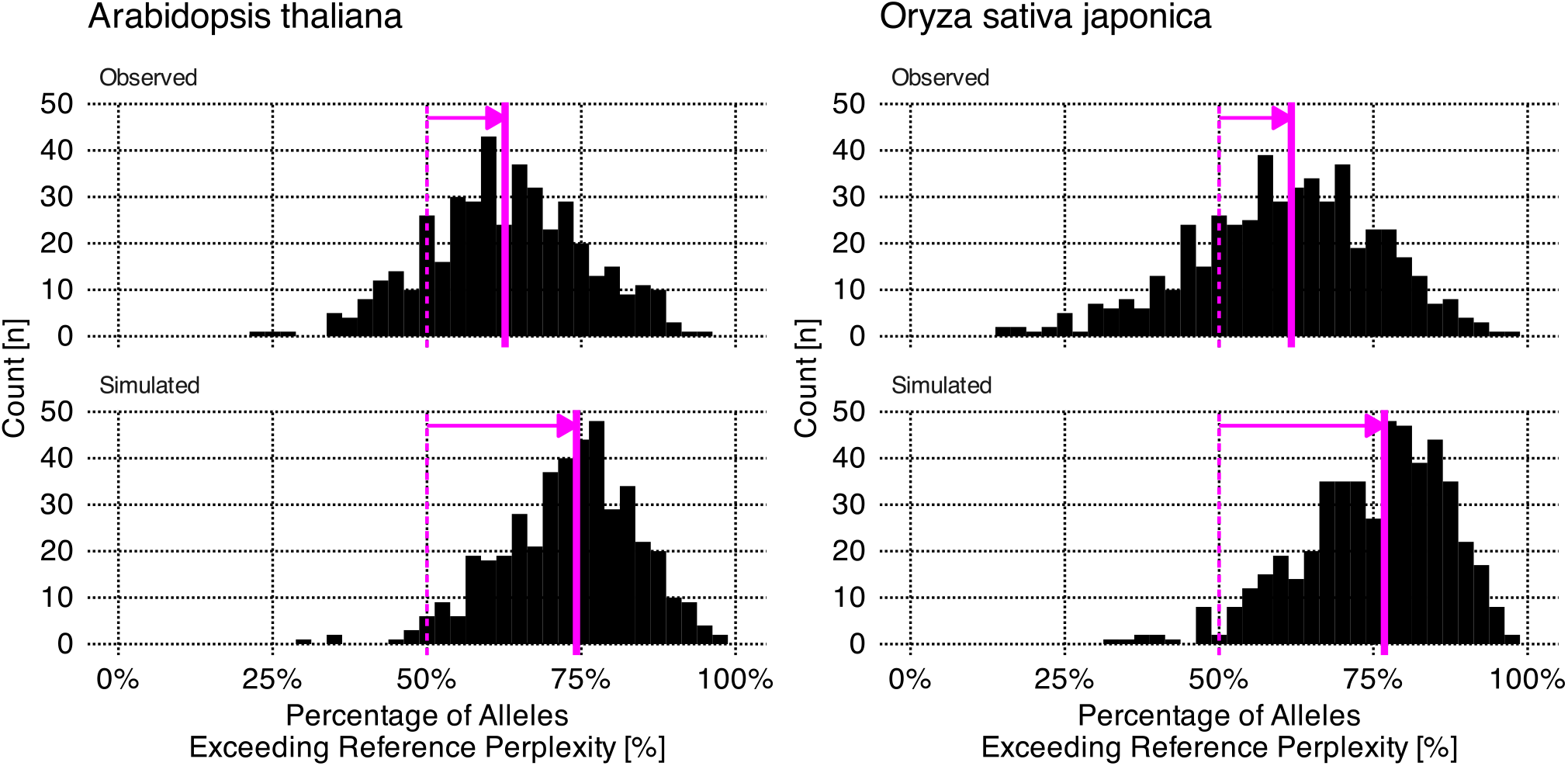
Perplexity distributions of single-nucleotide substitution alleles, relative to the perplexity of their respective reference allele. Each histogram data point represents one single gene, plotted as the percentage of its single-nucleotide substitution alleles with perplexity exceeding the one of their unmodified reference sequence. Results are shown stratified by species (*Arabidopsis thaliana* and *Oryza sativa japonica*) and by the nature of mutations (observed and simulated). The vertical thin dashed magenta line represents the expected center of the distribution, if single-nucleotide substitutions were equally likely to increase or decrease the allele’s perplexity; while the vertical solid magenta line represents the actual center of the distribution, measured as the median. The distinct shift of the distribution centers toward higher values indicates that mutated alleles typically possess higher perplexity than their reference counterparts, a trend more pronounced for simulated mutations.

We interpret this shift—which is stronger for simulated mutations—as evidence that Evo 2 is aligned with our empirical understanding of DNA sequences: it tends to assign lower probabilities to sequence variants that have not been recorded in living organisms and, therefore, might be less likely to occur. The fact that sequences with observed mutation tend to be perceived, by Evo 2, as more unlikely than their reference allele, is less straightforward to explain. One possibility is that many of these observed variants might be rare in natural populations, reinforcing the hypothesis that Evo 2 is aligned with our knowledge of plant genomes. An alternative explanation is that Evo 2 might be partially over-fitted to reference genome sequences. Further data collection on natural variation and on fitness of natural alleles might be needed to discriminate between these hypotheses.

### 3.7 Simulated mutations induce higher perplexity than observed mutations

In Section 3.6, we observed that the perplexity of simulated alleles tends to be higher than that of their observed counterparts within the same gene. To investigate this aspect in greater detail, we separately analyzed the perplexity distribution of observed and simulated alleles, further stratifying data into the four functional classes of mutations for each gene: (i) non-coding, (ii) silent, (iii) missense, and (iv) nonsense. We tested the statistical significance of our observations, the results of which are reported in Figure 7.

**Figure 7:**
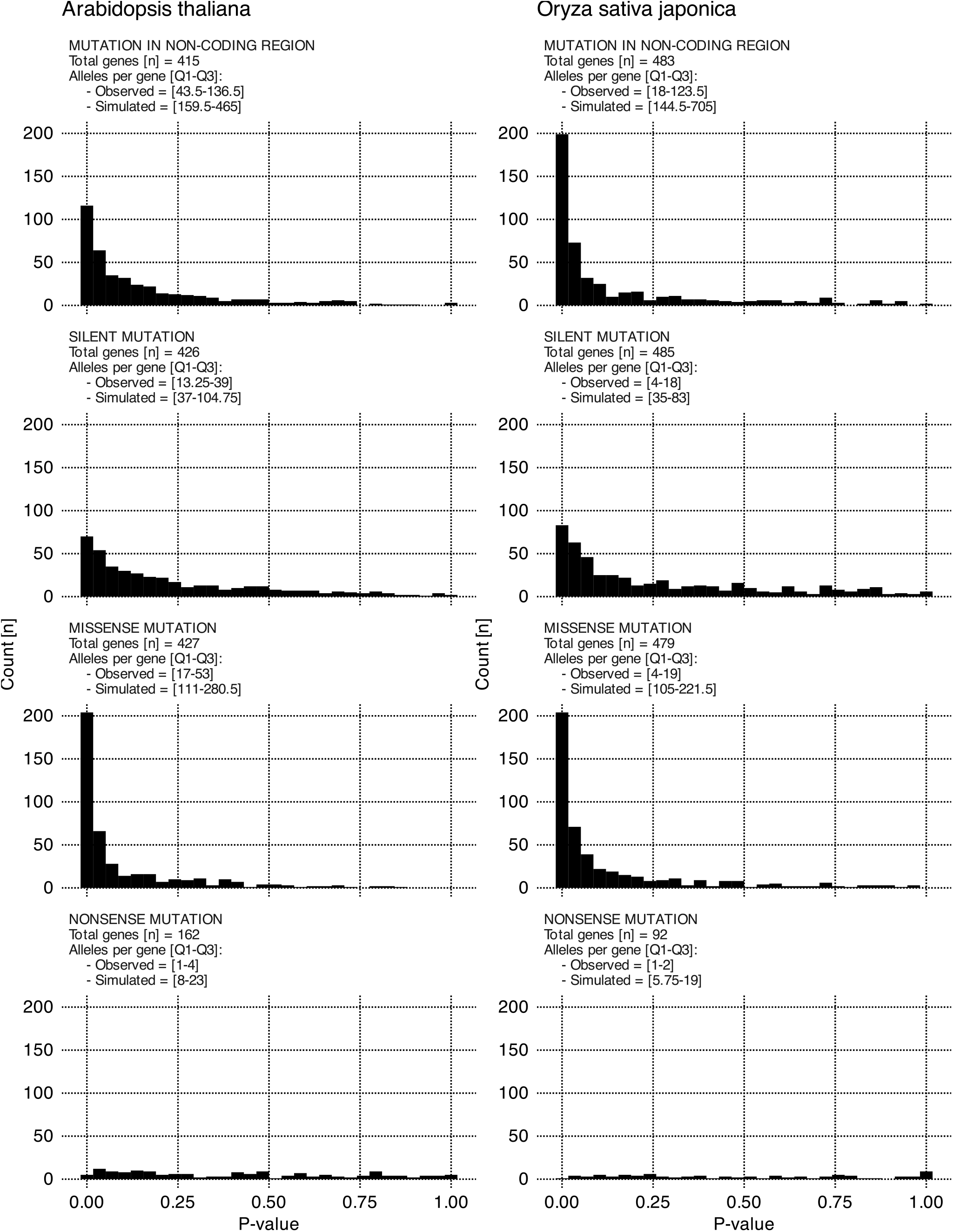
Distribution of p-values for one-sided Wilcoxon-Mann-Whitney test, comparing perplexity of observed and simulated alleles within each gene. Each data point in the histograms represents a gene-specific p-value. The alternative hypothesis stipulates that, for a given gene, alleles with a simulated mutation yield a higher perplexity than alleles with an observed mutation. Comparisons are stratified by species and mutation type. For each histogram we also report: (i) the total number of genes included in the comparison, and (ii) the interquartile range (IQR) of the number of substituted alleles per gene: this range depends on gene length for simulated mutation, and on available data for observed mutations. Most distributions are heavily right-skewed, indicating that for the majority of genes, the null hypothesis can be rejected. Across both species, we observe a strong effect for missense and non-coding mutations, a moderate effect for silent mutations, and no detectable effect for nonsense mutation.

The histograms in Figure 7 report the p-value distribution for one-sided Wilcoxon-Mann-Whitney tests across all genes considered in this study. The alternative hypothesis states that perplexity for simulated alleles is higher. While under the null hypothesis we would expect a uniform distribution of p-values between 0 and 1, we instead observe a strong right skew. This indicates that for the vast majority of genes, and mutation classes in both species, the null hypothesis can be rejected in favor of the alternative.

While the effect is statistically significant, the perplexity distributions of simulated and observed alleles tend to overlap extensively. This is quantified by the probability of superiority (Figure 8), defined as the empirical probability that a randomly sampled simulated allele has a higher perplexity than a randomly sampled observed allele from the same group.

**Figure 8:**
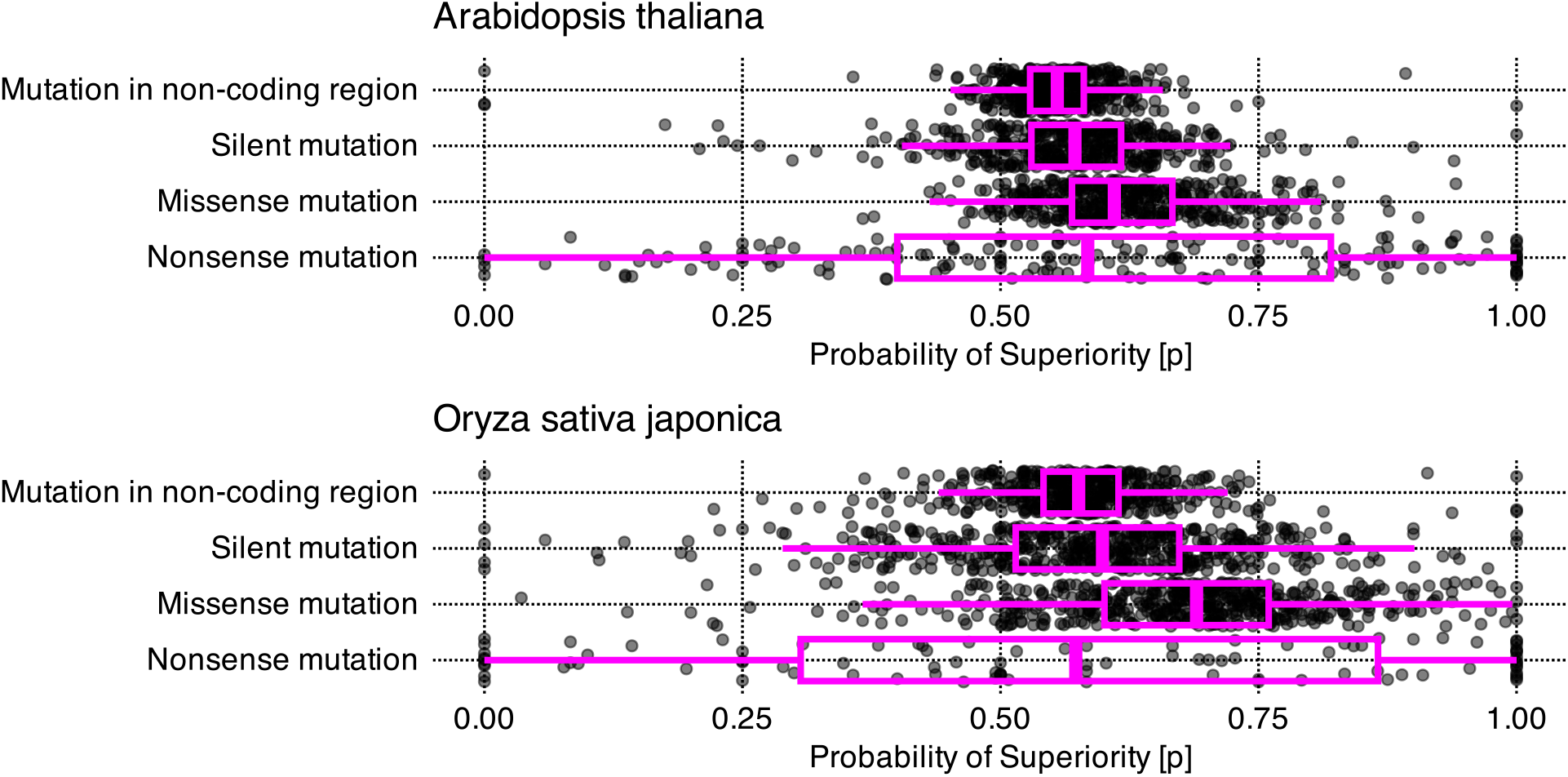
Distribution of probability of superiority comparing perplexity of observed and simulated alleles within each gene. Each data point represents a gene-specific probability of superiority for the perplexity of simulated alleles compared to that of observed alleles. Comparisons are stratified by species and mutation type. The probability of superiority represents the empirical probability that a randomly sampled simulated allele has a higher perplexity than an observed allele randomly sampled within the same group. In both species, we observed higher median probability of superiority for simulated missense mutations compared to their observed counterparts, followed by silent and non-coding mutations. Nonsense mutations probabilities of superiority are highly spread, possibly because of low sample numbers.

In genes from *Arabidopsis thaliana*, the median probabilities of superiority are 0.56, 0.57 and 0.61 for non-coding, silent and missense mutation. Results for *Oryza sativa japonica* follow a similar trend, with slightly higher scores: 0.58, 0.60 and 0.69 respectively. In contrast, nonsense mutations in both species showed highly dispersed probabilities, likely because small sample sizes constrained the precision of these measurements.

We interpret the statistically significant increase in perplexity yielded by simulated single-nucleotide substitution alleles, when compared to observed ones (Figure 7), as evidence that Evo 2 7B is indeed aligned with our current understanding of genetics constraints. On average, the model interprets simulated mutations as more unlikely than the ones observed in living organisms.

The only exception is the nonsense mutation class, where no significant effect was detected. We attribute this to a highly restricted sample size, which precludes inferring a distribution from data; this is a consequence of structural constraints on the number of nonsense variants that can be generated with one single-nucleotide substitution.

Nevertheless, the relatively modest probabilities of superiority (Figure 8), indicates that the distributions are often substantially overlapped. In most cases, simulated and observed alleles cannot be reliably distinguished from one another by perplexity alone. Future research could investigate whether using progressively shorter sequences, centered on the mutation site, enhances the resolving power of the model.

### 3.8 Single nucleotide substitutions affect Evo 2′s understanding of the local adjacent genetic sequences

Finally, we quantified the average magnitude of the perturbation induced by single-nucleotide substitutions on Evo 2′s next-nucleotide prediction probabilities, downstream of the mutation. We measured the distance required for this perturbation to decay, with prediction probabilities returning to a level comparable to that estimated on the unmutated reference. This analysis focused exclusively on missense mutations, as they exhibit the most robust and conserved effect across both species.

The magnitude and spatial decay of the effect of single-nucleotide substitutions—spanning the mutation site and the 25 nucleotides immediately downstream— are illustrated in Figure 9. The distribution of differences in the probabilities assigned by Evo 2 for the correct next nucleotide (between substituted and reference alleles) is strongly shifted toward negative values. This indicates that the mutated sequences receive lower probabilities than the reference at, both, the substitution site and subsequent downstream positions. Notably, the effect is more pronounced for simulated mutations than for the observed ones, in agreement with results reported in Section 3.6 and in Section 3.7. The resulting perturbation profiles appear to follow an exponential decay as the distance from the mutation site increases (Figure 9).

**Figure 9:**
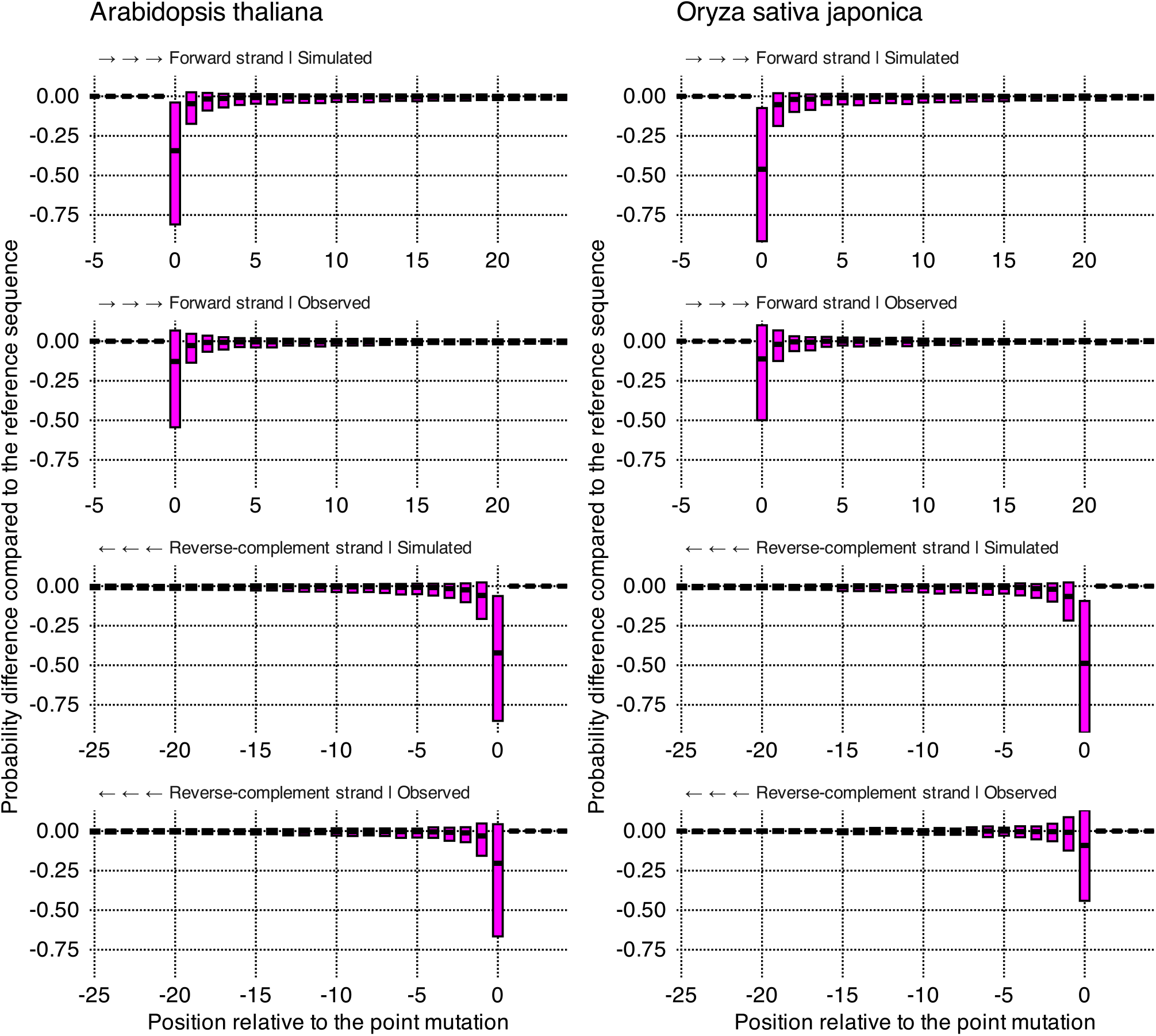
Magnitude and decay of the effect of single-nucleotide missense substitutions on Evo 2′s interpretation of the adjacent genetic sequence. Each data point used to draw the boxplot corresponds to a difference in the probability of Evo 2 predicting the next nucleotide correctly between a single-nucleotide substituted allele and the reference allele. This analysis was carried out on a pseudorandom 10% subsample of the genes examined in this study. Distances on the x-axis are expressed relative to the mutated nucleotide (position 0) in the 5′ to 3′ direction of the forward strand. The boxplots represent medians and interquartile ranges; whisker/boundaries and outliers are omitted. For each species we show four conditions: (i) observed-mutation forward strand, (ii) simulated-mutation forward strand, (iii) observed-mutation reverse strand, and (iv) simulated-mutation reverse strand. A visual evaluation indicates that the effect of single-nucleotide substitutions is consistently larger in magnitude for simulated mutations in both species considered, both when scoring the forward strand and the reverse strand.

Further analysis of the decay of this effect (Table 2) reveals that the impact of a single-nucleotide substitution remains detectable for several hundred nucleotides downstream, occasionally extending beyond 1000 nucleotides. Notably, the effect appears to be more persistent for observed mutations, particularly in *Oryza sativa japonica* compared with *Arabidopsis thaliana*. The mechanism underlying this species-specific variation remains unknown and warrants further investigation.

## 4 Conclusions

The evaluation of foundation DNA-LLM capabilities is currently constrained by our incomplete understanding of the functional and evolutionary forces shaping genome sequences (the current state and challenges of DNA-LLMs are reviewed in Hassan et al. (2025) and Consens et al. (2025)). With this challenge in mind, we present an approach that utilizes single-nucleotide substitutions to study how Evo 2 7B, one of the most advanced publicly available foundation DNA-LLM, interprets gene sequences of two plant model organisms. Relying on random sampling and on two fundamental categories to classify mutations, we could minimize the assumptions made on the mutations under study.

Relying on descriptive statistics and on non-parametric tests, together with perplexity as a metric for predictive proficiency of the model over a whole sequence, we analyzed and quantified the model’s comprehension of gene sequences, when single nucleotides are substituted. In our findings, across both *Arabidopsis thaliana* and *Oryza sativa japonica*, alleles sequences containing simulated single-nucleotide substitutions that, according to our current knowledge of functional and evolutionary genomics, might be more likely to disturb the gene’s function, are assigned on average, higher perplexity (i.e., lower prediction probabilities). In detail, Evo 2 7B tends to penalize simulated alleles compared to naturally occurring ones, mirroring our current understanding of genome evolution and its selective constraints.

In our results, while perplexity distributions of simulated and of observed alleles are statistically distinct, their significant overlap prevents the use of perplexity alone to identify the origin of randomly sampled alleles. This lack of clear separation is expected, as observed mutations are only a sample of those likely to be observed, and because the likelihood of a mutation to be observed is not dictated by the proximal sequence alone. Rather, it is also governed by a complex interplay of distal sequences, of endogenous factors—such as the targeting of DNA repair mechanisms mediated by epigenetic signal H3K4me1 (Quiroz et al., 2024), of cell-type-specific dynamics (Amundson et al., 2025), and of various exogenous biotic and abiotic influences, as reviewed by Quiroz et al. (2023).

Given this variability, capturing the true likelihood density of mutations may require models with multimodal inputs. While one could argue that Evo 2′s likelihoods represent evolutionary fitness rather than raw mutation probability, fitness itself is similarly contingent on endogenous and exogenous contexts the model lacks. Consequently, because DNA-LLMs rely solely on sequence input, the precise biological interpretation of next-nucleotide likelihoods and sequence perplexity remains an open question. Nevertheless, it is encouraging that Evo 2, even with a gene-sized context window, favors alleles observed *in vivo*. While we currently utilize perplexity estimated on a full gene sequence for its simplicity and standardization, future refinements to context windows and evaluation metrics may improve our ability to discriminate between observed and simulated mutations.

While single-nucleotide substitutions are included within the extensive evaluation strategies employed by Brixi et al. (2026) to assess Evo 2′s proficiency across human and multiple other eukaryotic genomes, and are included in the DART-Eval benchmark (Patel et al., 2024), our approach offers a distinct alternative. Although it decreases the number of species under study, our approach minimizes modeling assumptions through random sampling and increases the diversity of mutations evaluated.

Overall, DNA-LLMs are increasingly capable of extracting, representing, and predicting complex genetic features across diverse taxa (Shu et al., 2026). These models are already demonstrating utility in oncology (Zeng et al., 2025), infectious disease mitigation (Testagrose et al., 2025), and precision plant breeding (Liu et al., 2025). However, while results are promising, the sheer complexity of genomes, especially the eukaryotic ones, ensures that benchmarking foundation DNA-LLMs remains a significant challenge. We envision that future progress may require evaluating increasingly complex mutational landscapes and, crucially, transitioning toward *in vivo* validation. While recent literature reports successful *in vivo* validation efforts in prokaryotic systems, in generating *de novo* functional plasmids (Cunningham et al., 2025) and genes (Merchant et al., 2025), translating these workflows to higher organisms, such as plants, may exceed current experimental capabilities for assessing the *in vivo* effects of mutations.

## 5 Data and software availability

The substituted sequences and the collection of scripts necessary to reproduce this study are available at https://data.europa.eu/doi/10.2905/JRC.JH9JHTG. The scripts are mirrored at https://github.com/othomantegazza/sns-evo2.

## 6 Writing and editing disclosures

The authors utilized GPT-OSS 120b and Gemini 3 Flash exclusively to review and improve the readability of the English language employed in this study. Following these refinements, the final content was further reviewed and approved by the authors.

## 7 Competing interests

The authors declare no competing interests.

## 8 Grant information

This study was funded by the Exploratory Research Programme of the Joint Research Centre—European Commission.

## 9 Acknowledgements

The authors gratefully acknowledge Lia Orfei for her valuable advice on selecting suitable statistical approaches for this study, as well as Marco Mazzara, Koen Jonkers and Ursula Vincent for their support.

## Notes

https://data.europa.eu/doi/10.2905/JRC.JH9JHTG

### Competing Interest Statement

The authors have declared no competing interest.

### Summary of Updates

To make the code easier to access and download, a link to an external github repo that mirrors the original code repository was added to the manuscript.

